# Preparing for the worst: Evidence that older adults proactively downregulate negative affect

**DOI:** 10.1101/359901

**Authors:** Brittany Corbett, M. Natasha Rajah, Audrey Duarte

## Abstract

Age-related differences in processing emotional stimuli are well established. However, previous studies have only assessed the impact of age on emotional processing and encoding in response to, not in anticipation of, emotional stimuli. In the current study, we investigated age-related differences in the impact of emotional anticipation on affective responses and episodic memory for emotional images. Young and older were scanned while encoding negative and neutral images preceded by cues that were either valid or invalid predictors of image valence. Participants were asked to rate the emotional intensity of the images and to complete an episodic recognition task immediately after scanning. Using multivariate behavioral partial least squares (PLS) analysis, we found that young and older adults recruit the same set of brain regions to differentially support emotional processing during the anticipation of emotional images. Specifically, anticipatory recruitment of the amygdala, ventromedial PFC, and hippocampus in older adults predicts reduced memory for negative than neutral images for older adults and the opposite for young adults. Seed PLS analyses further show inverse coupling between the amygdala and ventromedial PFC activation following negative cues, consistent with the top-down spontaneous suppression of negative affect. To the best of our knowledge, this is the first study to provide evidence that the “positivity effect” seen in older adults’ memory performance is related to the spontaneous suppression of negative affect in anticipation of, not just in response to, negative stimuli.

## 1. Introduction

Though many cognitive abilities have been found to decline with age, emotional processing seems to improve with age. This has been described as both an age-related shift towards positive, and away from negative, information (Mather & Carstensen, 2005). For example, eye tracking studies have shown that older adults fixate more on neutral than negative images, whereas young adults show the opposite pattern (Isaacowitz, Wadlinger, Goren, & Wilson, 2006; Knight et al., 2007; Mather & Carstensen, 2003). In memory studies, older adults typically remember more positive and/or neutral than negative stimuli, whereas young adults remember more negative stimuli (for review, Mather, 2012). This “positivity effect” in older adults is best explained by the Socioemotional Selectivity Theory (SST), which states: as individuals age and their perceived time left in life diminishes, they focus more on improving their quality of life by regulating their emotions and less on other goals, such as information seeking (Carstensen, Isaacowitz, & Charles, 1999).

Consistent with SST, a number of studies have found that older adults are more likely to spontaneously downregulate negative affect than young adults. That is, when not given instructions to regulate, older adults spontaneously engage in emotional regulation strategies, whereas young adults have to be explicitly instructed to do so (for review, Mather, 2012). When older adults do spontaneously downregulate, self-reports suggest that they are more likely to engage in suppression than any other type of emotional regulation strategy (Eldesouky & English, 2018; Nolen-Hoeksema & Aldao, 2011). Suppression is defined as inhibiting one’s outward emotional reaction (for review, Richards, 2004). During negative reappraisal, individuals try to reinterpret the emotional meaning of the stimulus. This elaborative processing during reappraisal tends to result in enhanced memory for the emotional stimulus, but this avoidance during suppression tends to result in impaired memory (Dillon, Ritchey, Johnson, & LaBar, 2007). Consistent with self-reports, suppression seems to be the most likely explanation for the pattern of results we see in older adults outlined above, notably the negativity avoidance and impaired memory for negative events.

The ‘cognitive control model’ extension of the SST posits that the ability to regulate emotion is dependent upon prefrontal cortical (PFC) supported cognitive control processes (Mather & Knight, 2005). Older adults with higher cognitive control abilities, as determined with neuropsychological tests, are more likely to show the positivity effect than those with lower cognitive control abilities (Petrican, Moscovitch, & Schimmack, 2008). This theory is also supported by a number of neuroimaging studies that have found an increase in medial PFC activity for negative compared to neutral stimuli in older, but not young adults (for review, Nashiro, Sakaki, & Mather, 2012). Moreover, this increase in medial PFC activity, particularly the ventromedial PFC (vmPFC) is coupled with a decrease in amygdala activity during viewing of negative images (Gunning-Dixon et al., 2003; Leclerc & Kensinger, 2011; Sakaki, Nga, & Mather, 2013; St Jacques, Dolcos, & Cabeza, 2010; Urry et al., 2006). This inverse relationship is proposed to be reflective of successful emotional regulation, such that the increased vmPFC activity represents older adults’ employment of cognitive control strategies to reduce negative affect, which is mediated by amygdala activity (Leclerc & Kensinger, 2011).

Previous studies have only assessed the impact of age on emotional processing and encoding in response to emotional stimuli. However, in the real world, we can often anticipate emotional events, such as a scheduled surgery or screeching tires before an accident, which allows us to prepare for and regulate our emotional reaction. This leads one to question if age-related differences in emotional processing and memory are due solely to changes in the recruitment of processes elicited by stimuli or also in those recruited in anticipation of these stimuli. The impact of emotional anticipation on selected attention, memory and brain activity has been well-investigated in young adults (Galli, Wolpe, & Otten, 2011; Grupe, Oathes, & Nitschke, 2013; Mackiewicz, Sarinopoulos, Cleven, & Nitschke, 2006; Nitschke, Sarinopoulos, Mackiewicz, Schaefer, & Davidson, 2006). As outlined above, even though aging is known to affect each of these aspects of emotional processing, virtually nothing is known about the impact of emotional anticipation on these processes in older adults. Emerging results from young adult studies suggest that the neural activity during the time-period preceding stimulus presentation is also sensitive to episodic memory performance. The majority of these studies have investigated ‘pre-stimulus subsequent memory effects’ during encoding (Cohen et al., 2015; Galli, Choy, & Otten, 2012; Gruber & Otten, 2010; Otten, Quayle, Akram, Ditewig, & Rugg, 2006; Otten, Quayle, & Puvaneswaran, 2010; Padovani, Koenig, Brandeis, & Perrig, 2011). Although the mechanisms underlying these effects are not entirely clear, existing evidence suggests that they are highly dependent on the same factors that influence encoding and retrieval effects like stimulus modality, emotional valence, type of encoding strategy implemented, reward incentives, and the format in which memory is tested. Consequently, pre-stimulus memory effects are thought to reflect, at least in part, preparatory mobilization of processes that contribute to memory performance (Adcock, Thangavel, Whitfield-Gabrieli, Knutson, & Gabrieli, 2006; Addante, de Chastelaine, & Rugg, 2015; Otten et al., 2006). It is possible that at least some previous evidence of age-related differences in emotional memory encoding may be more accurately characterized as onsetting prior to stimulus presentation.

Typically, in emotional anticipation studies, young adults are given a cue (i.e. a X or O) that symbolizes the valence of the following image, thus allowing young adults to anticipate the upcoming image (Galli et al., 2011; Grupe et al., 2013; Mackiewicz et al., 2006; Nitschke et al., 2006). These studies found that the same brain regions involved in the presentation of negative images are also activated during the anticipation of negative images. These regions include a number of emotional processing and cognitive control regions: the amygdala, the anterior cingulate cortex (ACC), the medial PFC (mPFC), the dorsolateral PFC (DLPFC), the insula and the hippocampus (Mackiewicz et al., 2006; Nitschke et al., 2006; Simmons, Matthews, Stein, & Paulus, 2004). Interestingly, Mackiewicz et al. (2006) found that the anticipatory activity in the amygdala and hippocampus predicted successful memory encoding of negative stimuli above and beyond picture viewing activity, suggesting that emotional anticipation plays a crucial role in attention and cognitive processes that likely lead to elaborative encoding of the following stimuli in young adults.

However, there is often uncertainty regarding the emotional significance of future events. For example, one might be asked to unexpectedly meet with their boss and given recent layoffs in the company, one may start to anticipate that they are getting laid off in that meeting. This uncertainty reduces one’s ability to effectively prepare an optimal response and potentially enhance anxiety (Grupe & Nitschke, 2011). Generally, the effect of uncertainty is investigated by comparing certain negative events with uncertain negative events. The findings of these studies indicate that uncertain events are associated with stronger affective responses through self-reported mood, anxiety, and valence ratings (Bar-Anan, Wilson, & Gilbert, 2009; Grupe & Nitschke, 2011; Nader & Balleine, 2007; Yoshida, Seymour, Koltzenburg, & Dolan, 2013), physiological reactivity and hypervigilance (Grillon, Baas, Lissek, Smith, & Milstein, 2004; Nelson, Hodges, Hajcak, & Shankman, 2015), and greater activity in the amygdala, insula, ACC and lateral PFC (Ploghaus et al., 2001; Sarinopoulos et al., 2010; Yoshida et al., 2013). In sum, these findings point to an uncertainty-dependent defensive state (i.e. threat-related vigilance) employed by young adults that results in enhanced attentional processing and emotional salience detection, likely aiming at threat elimination.

The current study investigated age-related differences in the impact of emotional uncertainty on the anticipatory processes engaged prior to encoding of naturalistic, negative and neutral events. We manipulated participants’ emotional expectations using pre-stimulus cues in which a proportion invalidly indicated the valence of the impending image. To the best of our knowledge, this is the first fMRI study to investigate age-related differences in the impact of emotional anticipation on affective responses and memory for emotional events. We utilized behavioral partial least squares (PLS) analysis to determine the relationship between anticipatory neural activity, subjective emotional response ratings, and memory in young and older adults (McIntosh & N. J. Lobaugh, 2004). Additionally, we utilized seed PLS to assess the functional connectivity among the regions revealed in our behavioral PLS. Based on previous findings, we hypothesize that older adults, but not young adults, will engage in spontaneous downregulation of negative affect following negative cues. Behaviorally, this will result in lower emotional intensity ratings and memory performance for negative valid than negative invalid events. Cue-related activity should be greater following negative cues than neutral cues in regions implicated in emotional processing (amygdala, medial PFC) and further, we should see evidence of an inverse coupling between the vmPFC and amygdala, consistent with successful downregulation.

## 2. Method

### 2.1 Participants

Group Characteristics are presented in **Table 1.** The participants for this study were 22 young adults (13 females, ages 18-38, mean age = 22.68) and 20 older adults (10 females, ages 60-76, mean age = 67.6). Older adults were slightly more educated than young adults [*t*(40) = 2.022, p = .050]. Four additional young adults’ data were not included in subsequent analyses due to three participants’ responses not being recorded as a result of technical issues with our experimental program and one who terminated the experiment early. Two older adults’ data were excluded as well due to excessive motion during scanning and an additional two participants did not complete the experimental task. All participants were right-handed, native English speakers, with normal or corrected to normal vision (using MRI-compatible glasses when necessary). Participants were excluded from the study if they reported any of the following conditions: Epilepsy, Parkinson’s disease, a history of stroke or seizure, untreated depression, untreated anxiety, Attention Deficit Disorder, Multiple Sclerosis, uncontrolled hyper- or hypo-tension, untreated Diabetes, Sickle Cell Anemia, smoking or other regular use of nicotine, use of beta blockers, alcoholism and regular use of illegal drugs. All participants were recruited from Georgia Tech or the surrounding Atlanta community. Participants were compensated $10 an hour plus an additional $5 for travel expenses. Any young adults that were enrolled in a psychology course at Georgia Tech were given the option to receive extra credit instead of monetary compensation. All participants signed consent forms approved by the Georgia Institute of Technology Institutional Review Board.

**Table 1.**
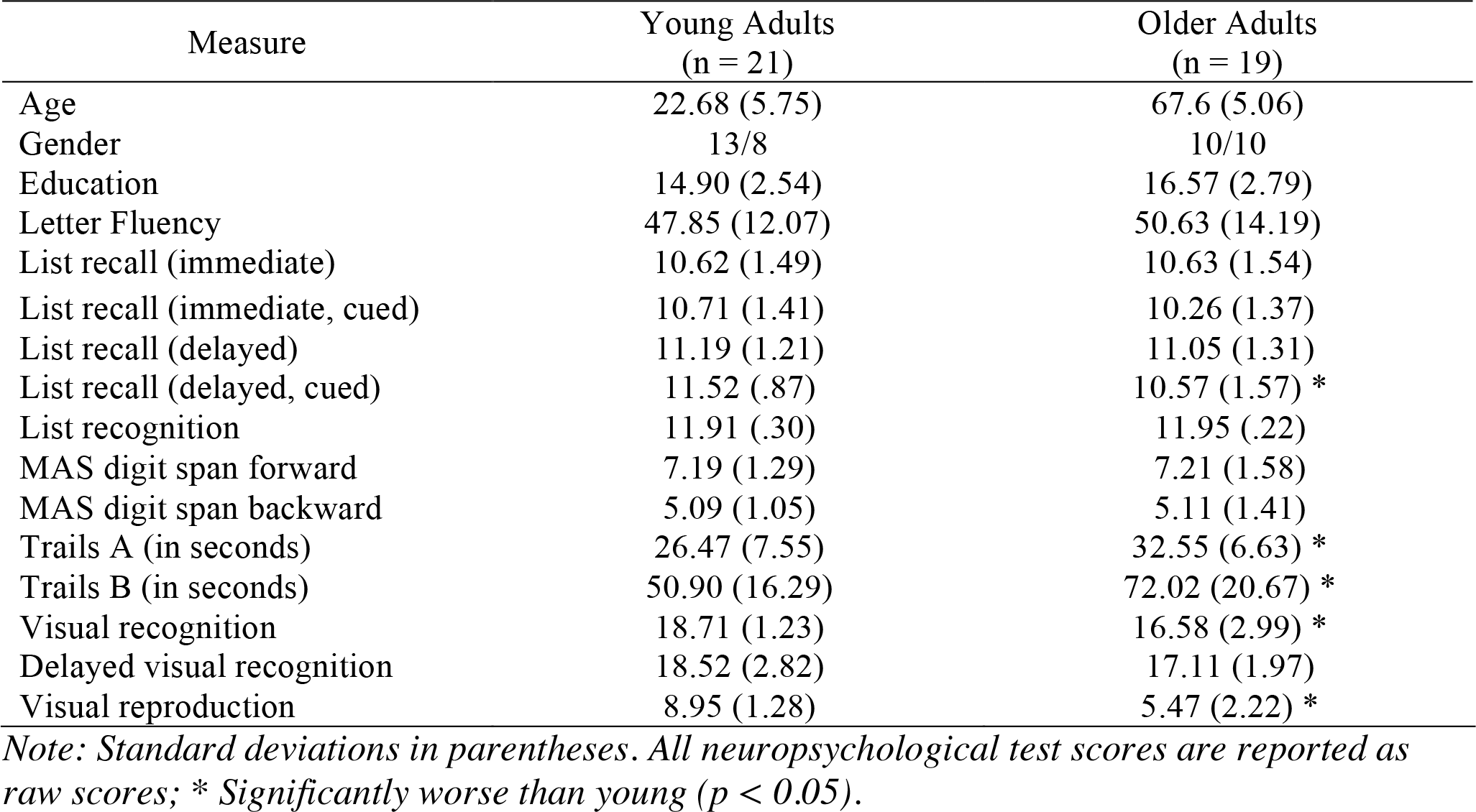
Group Characteristics

| Measure | Young Adults<br>(n = 21) | Older Adults<br>(n = 19) |
| --- | --- | --- |
| Age | 22.68 (5.75) | 67.6 (5.06) |
| Gender | 13/8 | 10/10 |
| Education | 14.90 (2.54) | 16.57 (2.79) |
| Letter Fluency | 47.85 (12.07) | 50.63 (14.19) |
| List recall (immediate) | 10.62 (1.49) | 10.63 (1.54) |
| List recall (immediate, cued) | 10.71 (1.41) | 10.26 (1.37) |
| List recall (delayed) | 11.19 (1.21) | 11.05 (1.31) |
| List recall (delayed, cued) | 11.52 (.87) | 10.57 (1.57) * |
| List recognition | 11.91 (.30) | 11.95 (.22) |
| MAS digit span forward | 7.19 (1.29) | 7.21 (1.58) |
| MAS digit span backward | 5.09 (1.05) | 5.11 (1.41) |
| Trails A (in seconds) | 26.47 (7.55) | 32.55 (6.63) * |
| Trails B (in seconds) | 50.90 (16.29) | 72.02 (20.67) * |
| Visual recognition | 18.71 (1.23) | 16.58 (2.99) * |
| Delayed visual recognition | 18.52 (2.82) | 17.11 (1.97) |
| Visual reproduction | 8.95 (1.28) | 5.47 (2.22) * |

*Note: Standard deviations in parentheses. All neuropsychological test scores are reported as raw scores; \* Significantly worse than young ( $p < 0.05$ ).*

### 2.2 Neuropsychological Assessment

Immediately after the recognition session, the participants were given a series of neuropsychological tests to ensure that no individual differences in task performance were due to cognitive impairments. The testing contained the Wechsler Adult Intelligence Test (WAIS-R) (Wechsler, 1981) digit-symbol substitution and digit span forward and backward tasks, the Shipley Vocabulary Test (Shipley, 1946) and the 64-card version of the Wisconsin Card Sorting Test (WCST) (Lezak, 1995). Any participant whose score fell 2 Standard-Deviations below their age-adjusted mean score given their level of education was removed from the sample. One older adult who had recently completed the assessment for another experiment and one young adult who was already familiar with test were excluded from testing.

### 2.3 Materials

Stimuli consisted of 480 colored images, 240 from the Nencki Affective Picture System (NAPS)(Marchewka, Zurawski, Jednorog, & Grabowska, 2013) and 240 from the International Affective Picture System (IAPS) (Lang, Bradley, & Cuthbert, 2008). We used both sets of images in order to achieve a sufficient number of highly arousing negative images. The images were selected to be the most unpleasant and arousing negative images based on published norms using the Self-Assessment Manikan (SAM) Scale, each ranging from 1 to 9 (1 = very negative, 9 = very positive; 1 = relaxed, 9 = aroused). In total there were 240 negative images (mean valence = 3.11, mean arousal = 6.07) and 240 neutral images (mean valence = 6.21, mean arousal = 4.01). The two auditory cues were chosen were from the International Affective Digital Sounds (IADS) (Bradley & Lang, 2007). The negative cue was a 1s clip of tires screeching (mean valence = 3.11, mean arousal = 6.07) and the neutral cue was a 1s clip of a wind chime (mean valence = 6.10, mean arousal = 4.23).

### 2.4 Procedure

A practice session was administered before each task to familiarize the participants with the task and to ensure they could perform the task. Only the encoding blocks were scanned. Stimuli were counterbalanced across participants such that each picture was not in the same category (valid vs invalid, old vs new) across participants. There were 320 stimuli shown during encoding, as well as an additional 160 stimuli shown during retrieval.

The encoding task consisted of 5 blocks, with 80 trials in each block. Each block consisted of 64 regular trials, with 80% of each of the trials preceded by a valid cue and 20% percent by an invalid cue, and 16 catch trials. The 80/20 valid/invalid ratio was used to ensure that participants did not distrust the cues (Wheeler et al., 2006). As seen in **Figure 1a**, each trial began with a central fixation cross for 300ms. Next, an auditory cue indicated either the correct (valid) or incorrect (invalid) valence of the upcoming picture, followed by a fixation cross for a cue stimulus interval (CSI) of 1000ms. The stimulus was then presented and the participant was asked to rate the emotional intensity of the image on a scale of 1-4 (1 being least intense – 4 being most intense). Participants were asked to make their responses on two button boxes, using only their index and middle fingers. They were asked to respond with their left middle finger as “1” for least intense, their left index finger as “2” for less intense, their right index finger as “3” for more intense and their right middle finger as “4” for most intense. A fixation cross was then presented on the screen for a fixed 500ms interval before the participant was asked to complete the arrows task. The arrows task maximizes design efficiency by pseudorandomly interspersing event trials with “active” baseline trials lasting between 2 and 6 s, jittered in increments of 2 s (Dale, 1999). Every 2 s, an arrow appears on the screen and participants are asked to respond using the button box to indicate the direction of the arrow: “1” in response to a left pointing arrow and “2” for a right pointing arrow. Requiring participants to respond to the arrows kept them engaged in the task and minimized default mode network activity (Stark & Squire, 2001). Catch trials differed from full trials in that no stimuli were presented following the cue. These trials were included to isolate cue-related activity from stimulus-related activity, as has been done in prior studies (Wheeler, M. E., Shulman, G. L., Buckner, R. L., Miezin, F. M., Velanova, K., & Petersen, S. E., 2006). Stimuli were presented in a pseudorandomized order in each block. Each block lasted 12 minutes and a brief rest period was allowed between each block to prevent fatigue. In total, the encoding session lasted 1 hour and 15 minutes.

**Fig. 1.**
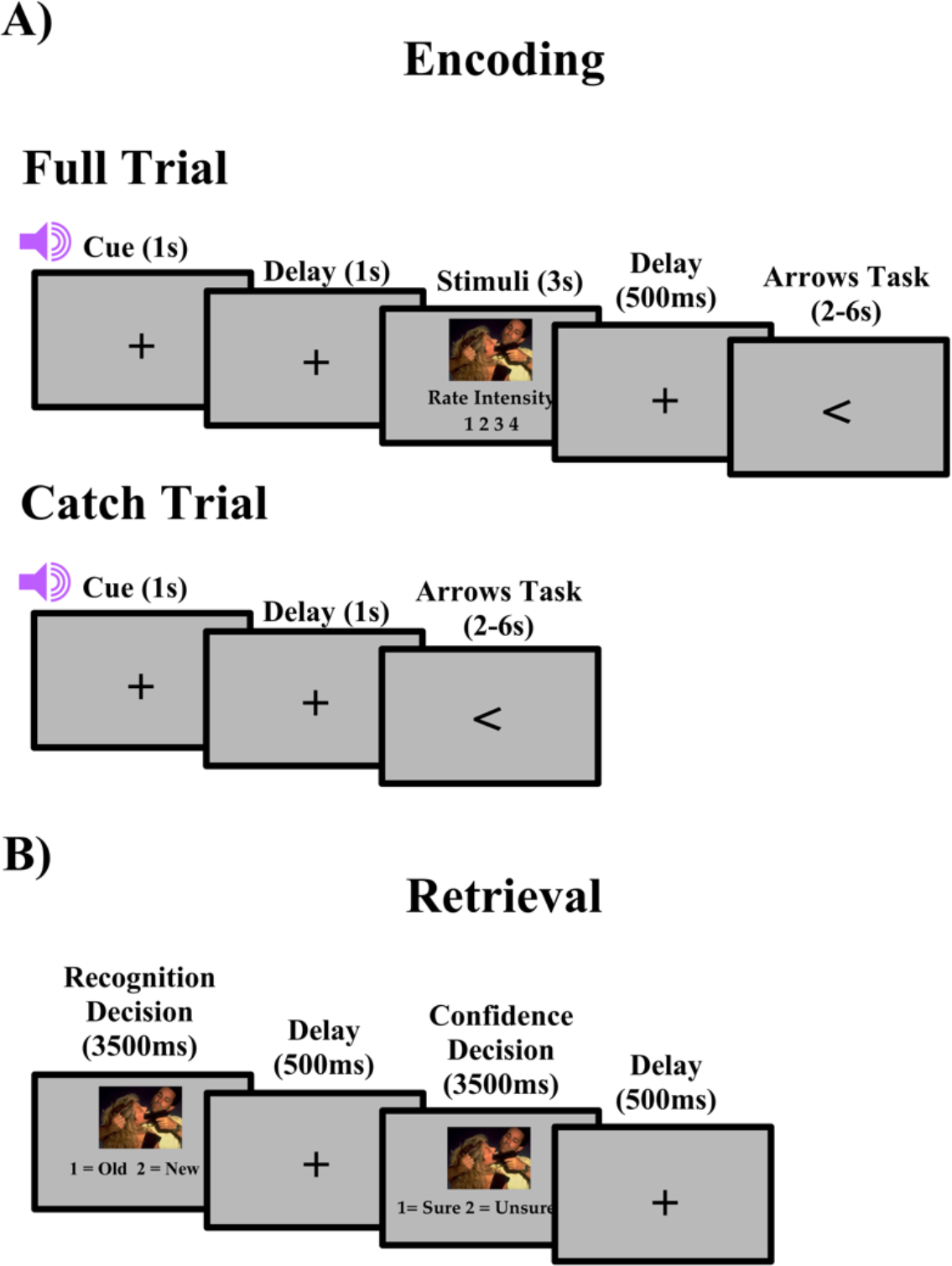
Task design for both encoding and retrieval. A) Full and catch trial design for encoding. B) Trial design for retrieval.

After the encoding task, participants immediately exited the scanner and began the recognition task. The recognition portion of the study was divided into 5 blocks with 96 trials, 64 old stimuli and 32 new stimuli in each block. For each trial, participants were asked to answer two questions about the picture (**Figure 1b**). First, participants viewed the picture for 3500ms and indicated if the image was old by responding with “1” on a keypad or new by responding with “2” on a keypad. Second, participants viewed the image again for 3500ms and made a confidence judgment, if they were sure about their recognition decision by responding with “1” or unsure by responding with “2”. A fixation cross was displayed in the center of the screen for 500ms between the questions. Confidence judgments were used to separate strong recognition from guessing, with unsure responses being considered as guesses.

### 2.6 fMRI Acquisition

Scanning was performed on a 3-T Siemens TIM Trio system. Functional data were acquired using a gradient echo pulse sequence (37 transverse slices oriented along the anterior– posterior commissural axis with a 30-degree upward tilt to avoid the eyes, repetition time of 2 s, echo time of 30 ms, 3 x 3 x 3.5 mm voxels, 0.8-mm interslice gap). Five encoding blocks of 371 volumes each were acquired. The first 2 volumes of each block were discarded to allow for equilibration effects. A high-resolution T1-weighted magnetization-prepared rapid acquisition gradient echo (MPRAGE) image was collected for normalization.

### 2.7 fMRI Analyses

#### 2.7.1 Preprocessing

Data was analyzed using SPM12 (SPM12, http://www.fil.ion.ucl.ac.uk/spm/software/spm12/). Images were corrected for differences in slice timing acquisition using the middle slice of each volume as the reference, spatially realigned and resliced with respect to the first volume of the first block. Each participant’s MPRAGE scan was coregistered to the mean EPI image, produced from spatial realignment. Each coregistered structural scan was then segmented using the Diffeomorphic Anatomical Registration Through Exponentiated Lie algebra (DARTEL) SPM 12 toolbox (Ashburner, 2007). DARTEL is a suite of tools fully integrated with SPM 12, which the SPM 12 manual recommends over optimized normalization, to achieve sharper nonlinear registration, for intersubject alignment. This method also achieves better localization of fMRI activations in Montreal Neurological Institute (MNI) space. This method has been used successfully in several previous studies with various healthy and neurological populations (Pereira et al., 2010; Yassa & Stark, 2009). Briefly, the gray and white matter segmented images were used to create a study-specific template using the DARTEL toolbox and the flow fields containing the deformation parameters to this template for each subject were used to normalize each participant’s realigned and resliced EPIs to MNI space. Normalized EPI images will be written to 3 × 3 × 3 mm and smoothed with an 8 mm full-width at half-maximum isotropic Gaussian kernel.

#### 2.7.2 Multivariate partial least squared analysis

Multivariate behavioral PLS was used to analyze the fMRI data to identify whole-brain patterns of task-related activity at encoding that correlate with subsequent memory accuracy and/or subjective emotional intensity ratings in young and older adults (A. R. McIntosh & N. J. Lobaugh, 2004). We chose this approach because PLS is a powerful data-driven method that identifies distributed patterns of activated voxels that differ across experimental conditions and/or relate to a specific behavioral measure without using pre-specified contrasts, and does not require that the assumptions of normality, independence of observations and linearity for general linear models be met (Van Roon, Zakizadeh, & Chartier, 2014).

##### 2.7.2.1 Cue Period Behavioral PLS

As our primary interest was to investigate regions recruited during the anticipation of emotional events that correlate with subsequent emotional memory accuracy and subjective emotional intensity ratings, we only included catch trails (i.e. trials in which the participant was only presented a cue, but no stimulus) in this behavioral PLS. There were two conditions, negative catch trials and neutral catch trials, and two groups, young adults and older adults, in this analysis. fMRI data for these catch trials were stored in a data matrix that was organized by condition and stacked across participants, within age-group, and age-groups were stacked above one another as follows: young-negative catch trial, young-neutral catch trial, old-negative catch trial, and old-neutral catch trial. fMRI data for each event onset (time lag = 0) as well as the ensuing seven volumes (2 sec x 7 = 14 sec) following each event onset was included in the data matrix. We then created two behavioral vectors which were stacked in the same order as our fMRI data matrix. The first vector included each individual’s memory accuracy benefit for negative compared to neutral events calculated as d-prime for negative minus d-prime for neutral events. The second vector included each individual’s intensity rating difference between negative and neutral events. Because participants did not know if a picture was valid or invalid until stimulus presentation, we only investigated valence effects in this analysis and thus, our behavioral measures were collapsed across cue validity. The two behavioral vectors were cross-correlated with the fMRI data. Singular value decomposition (SVD) of the resulting cross-correlation matrix was conducted to yield a set of orthogonal latent variables (LVs). Each LV contains: 1) a singular value, which reflects the amount of covariance accounted for by the LV; 2) a singular image, showing a pattern of whole-brain activity that is symmetrically related to 3) a correlation profile, which depicts how subjects’ memory accuracy and intensity ratings related to the pattern of brain activity observed in the singular image. The singular image includes brain saliences, which are numerical weights assigned to each voxel at each TR/time lag included in the data matrix. Brain saliences can be negative or positive. Brain regions with positive voxel saliences are positively related to the correlations profile, and those with negative voxel saliences are negatively related to the correlational profile. Thus, the pattern of whole brain activity identified by the singular image is symmetrically related to the correlation profile.

The significance of LVs was assessed through permutation tests on the singular values using an alpha of *p* < .05 and 1000 permutations. The permutation test involved sampling each subjects’ behavioral measure and event-related activity without replacement to randomly reassign the behavior-brain activity associations. For each permuted iteration a PLS was recalculated, and the probability that the permuted singular values exceeded the observed singular value for a given LV was used to assess significance at *p* < 0.05 (McIntosh, Chau, & Protzner, 2004). The permutation method used met the exchangeability criterion as described in McIntosh and N. J. Lobaugh (2004). The standard error for each singular image was calculated by conducting 500 bootstraps to sample subjects with replacement, while maintaining the experimental and group order. The ratio of the original brain salience to the bootstrap standard error (bootstrap ratio (BSR) was used to identify maximal reliable patterns of positive and negative brain saliences represented by the singular image. In the current study, BSR significance was set to ± 3.28, which is equivalent to *p* < .001, with a minimum cluster size = 15. In order to determine the subset of time lags that maximally represented the correlation profiles of LVs, we computed temporal brain scores for each condition in each significant LV (see A. R. McIntosh & N. J. Lobaugh, 2004 for further details). Using these temporal brain scores, we identified the peak time lags in this B-PLS to be 2-6 (4-12s post-event onset). All activations reported for this analysis will only be within these lags.

##### 2.7.2.2 Cue Period Seed PLS

Seed PLS was conducted to assess the functional connectivity of regions identified in our behavioral PLS. Specifically, we wanted to assess the functional connectivity between the ventromedial PFC (vmPFC) and the amygdala to determine if older adults to a greater extent than young adults were engaging in spontaneous emotional regulation following the presentation of the cue. We identified one seed voxel in the right vmPFC [18 66 −3]. This seed was selected for two reasons: one, it was reliably activated during our behavioral PLS and two, it is well evidenced that this region is recruited during emotional processing and regulation (for review, Nashiro et al., 2012). This seed voxel was defined by extracting the average response for each participant for each lag using a neighborhood voxel of 0 around the seed voxel. All voxels included around the seed voxel met the bootstrap and spatial extent threshold criteria in the behavioral PLS above. Seed PLS was conducted identically to the behavioral PLS described above except that instead of using behavioral measures we used each subject’s BOLD signal value from the seed voxel averaged across lags 2-6 as the vector.

## 3. Results

For all behavioral statistical tests, the Huynh-Feldt correction was applied where appropriate and is reflected in the *p* values and the error terms.

### 3.1 Neuropsychological Assessment Results

Group characteristics are shown in **Table 1.** All participants scored within 2 standard-deviations of age-adjusted normative averages for all the neuropsychological tests. Older adults had significantly worse performance than young adults on several tests: Cued/Delayed List Recall, Trails A, Trails B, Visual Recognition, Delayed Visual Recognition and Visual Reproduction [*t*’*s* > 2.071, *p’s* < .045]. There were no other significant group differences [*t’s* < 1.372, *p’s* > .178].

### 3.2 Behavioral Results

#### 3.2.1 Emotional Intensity Ratings

We created an emotional intensity score by giving a 4 to highly intense emotional ratings, a 3 to moderately intense emotional ratings, a 2 to less intense emotional ratings, and a 1 to least intense emotional ratings. We then averaged all the rating scores for each stimulus category for each participant. The emotional intensity data is displayed in **Table 2**.

**Table 2.** Mean Emotional Intensity Ratings across Valence, Cue, and Age

|  |  | Young Adults | Older Adults |
| --- | --- | --- | --- |
| Neutral | Valid | 1.84(.51) | 1.93(.61) |
|  | Invalid | 1.82(.45) | 1.89(.55) |
| Negative | Valid | 2.76(.43) | 2.83(.46) |
|  | Invalid | 2.72(.43) | 2.63(.52) |

*Note: Standard Deviations in parentheses*

These data were submitted to a 2 Age (young, old) X 2 Valence (negative, neutral) X 2 Cue (valid, invalid) ANOVA. There was a main effect of Valence [*F*(1,40) = 148.493, *p* < .0001, η^2^ = .788], a main effect of Cue [*F*(1,40) = 7.164, *p* = .011, η^2^ = .152] and a Valence X Cue Interaction [*F*(1,40) = 5.146, *p* = .029, η^2^ = .114], all other effects were not significant [*F*’s < 1, *p*’s > .777, η^2^’s < .002]. Follow-up ANOVAs for each separate valence category revealed that the main effect of Cue was reliable for negative images [*F*(1,41) = 8.775, *p* = .005, η^2^ = .176], but not for neutral images [*F*(1,41) < 1, *p* = .400, η^2^ = .017]. Both young and older adults rated negative valid images as more intense than negative invalid images [*t*(41) = 2.962, *p* = .005].

#### 3.2.2 Memory Accuracy

Percentage of hits, misses, and false alarms for each of the stimulus categories and groups are shown in **Table 3**. Only high confidence responses were included in the analyses.

**Table 3.**
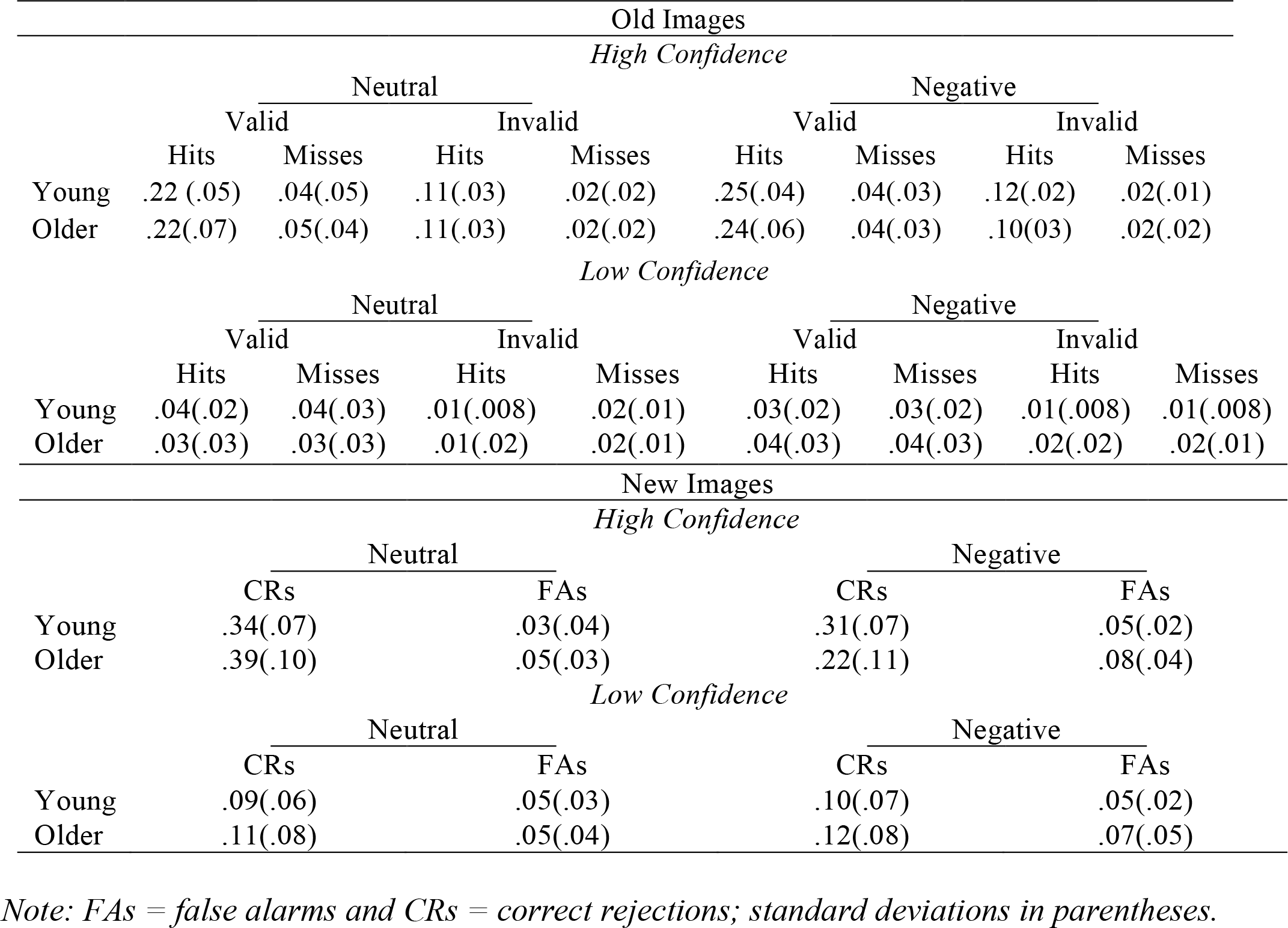
Mean Proportion of hits, misses, false alarms and correct rejections for young and older adults for each valence, validity and confidence category.

We analyzed high confidence item memory performance using signal detection theory [z(hit rate) – z(false alarm rate)]. The d’ data were submitted to a 2 Age (Young, Old) X 2 Valence (Negative, Neutral) X 2 Cue (Valid, Invalid) ANOVA. These data are presented in **Figure 3**. There was a main effect of Valence [*F*(1,40) = 14.170, *p* = .001, η^2^ = .262], a main effect of Age [*F*(1,40) = 21.764, *p* < .001, η^2^ = .352], a Valence x Age interaction [*F*(1,40) = 8.423, *p* = .006, η^2^ = .174] and a Cue x Age interaction [*F*(1,40) = 5.988, *p* = .019, η^2^ = .130], all other effects were not significant [*F*’s < 1, *p*’s > .311, η^2^’s < .017]. Follow-up ANOVAs for each age group showed that the main effect of Valence was reliable for older adults only. Older adults remembered more neutral than negative images [*F*(1,19) = 26.889, *p* < .001, η^2^ = .586], whereas young adults remembered a similar number of neutral and negative images [*F*(1,21) < 1, *p* = .573, η^2^ = .015]. The main effect of Cue was reliable for young adults only. Young adults remembered more invalid than valid images [*F*(1,21) = 5.179, *p* = .033, η^2^ = .198], whereas older adults remembered a similar number of invalid and valid images [*F*(1,19) = 1.327, *p* = .264, η^2^ = .065].

**Fig. 3.**
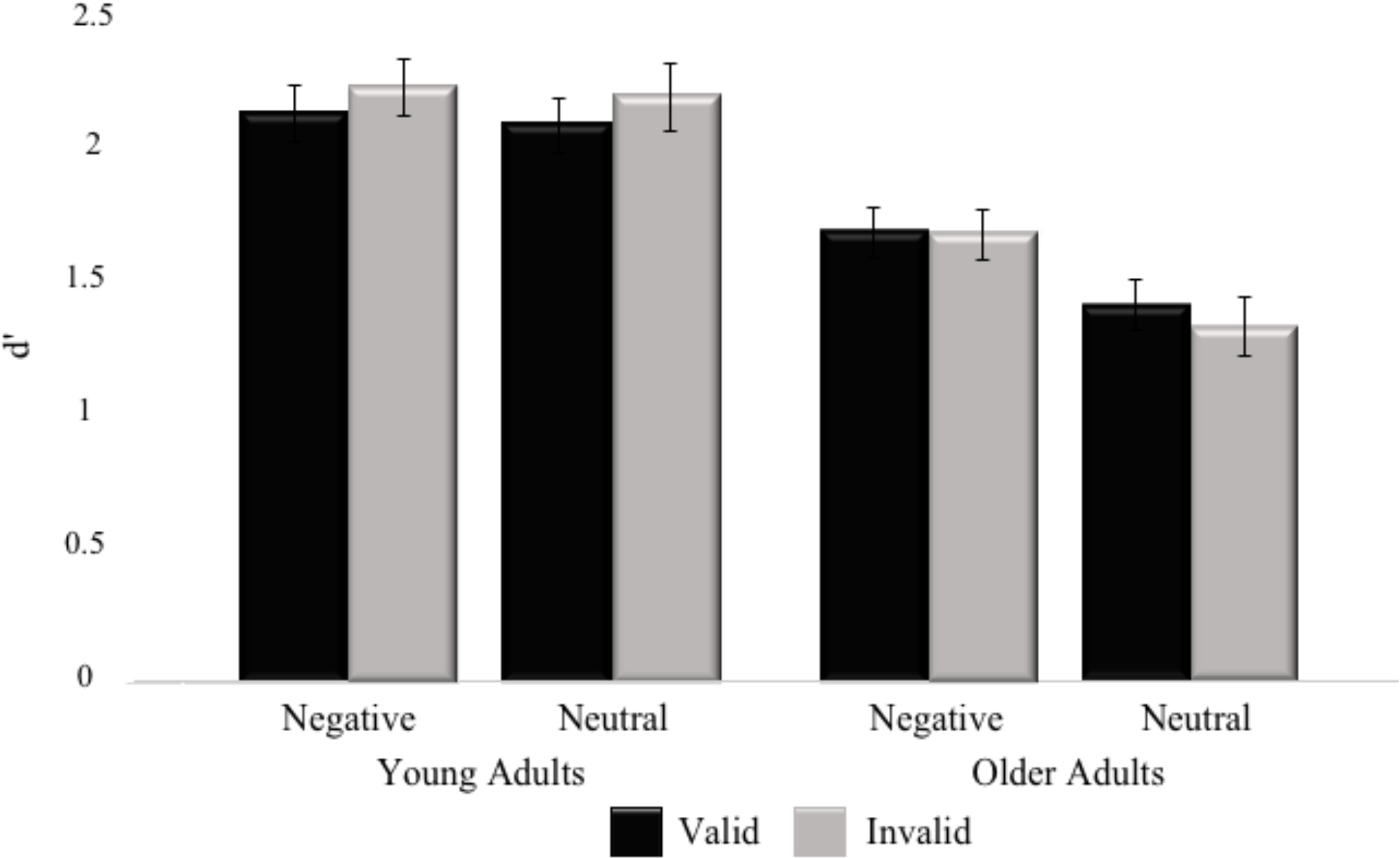
High Confidence memory accuracy (d’) for young and older adults for each stimulus category. Error bars represent the standard error of the mean.

We hypothesized that if older adults were engaging in spontaneous downregulation than intensity ratings would decrease with memory performance following negative, but not neutral cues. However, recent research has determined that when not given instructions on how to regulate, older adults are more likely to engage in suppression than any other type of emotional regulation strategy (Eldesouky & English, 2018; Nolen-Hoeksema & Aldao, 2011). As suppression has been found to impair memory performance, but have no effect on intensity ratings (Gross, 2002), we wanted to determine if both intensity ratings and memory accuracy were indicative of regulation in the present study. In order to determine this, we ran Pearson correlations between intensity ratings and memory accuracy for both valence categories. Results indicate that intensity ratings and memory accuracy were not correlated for young adults for both negative [*r*(22) = -.290, *p* = .191] and neutral images [*r*(22) = .004, *p* = .986], nor for older adults for both negative [*r*(20) = -.057, *p* = .811] and neutral images [*r*(20) = -.093, *p* = .696].

### 3.3 fMRI Results

#### 3.3.1 Cue Period Behavioral PLS

The behavioral PLS analysis identified one significant LV. This LV, which accounted for 31.49% of the total cross-block covariance (*p* < 0.001) is presented in **Figure 4a**, and the local maxima for this LV are presented in **Table 4**. Only negative salience brain regions from this LV withstood our spatial extent cutoff of a minimum of 15 clusters and BSR threshold of ± 3.28. The PLS correlation profile in **Figure 4b** demonstrates that during negative catch trials, activity in these negative salience regions increased with negative intensity ratings and with negative memory accuracy, relative to neutral intensity and accuracy, respectively for young adults but decreased with negative memory accuracy for older adults. During neutral catch trials, activity in these negative salience regions increased with negative intensity ratings for both young and older adults. These regions included: bilateral amygdala, bilateral anterior cingulate cortex, bilateral dorsomedial prefrontal cortex, right ventromedial prefrontal cortex, bilateral insula and the left hippocampus.

**Fig. 4.**
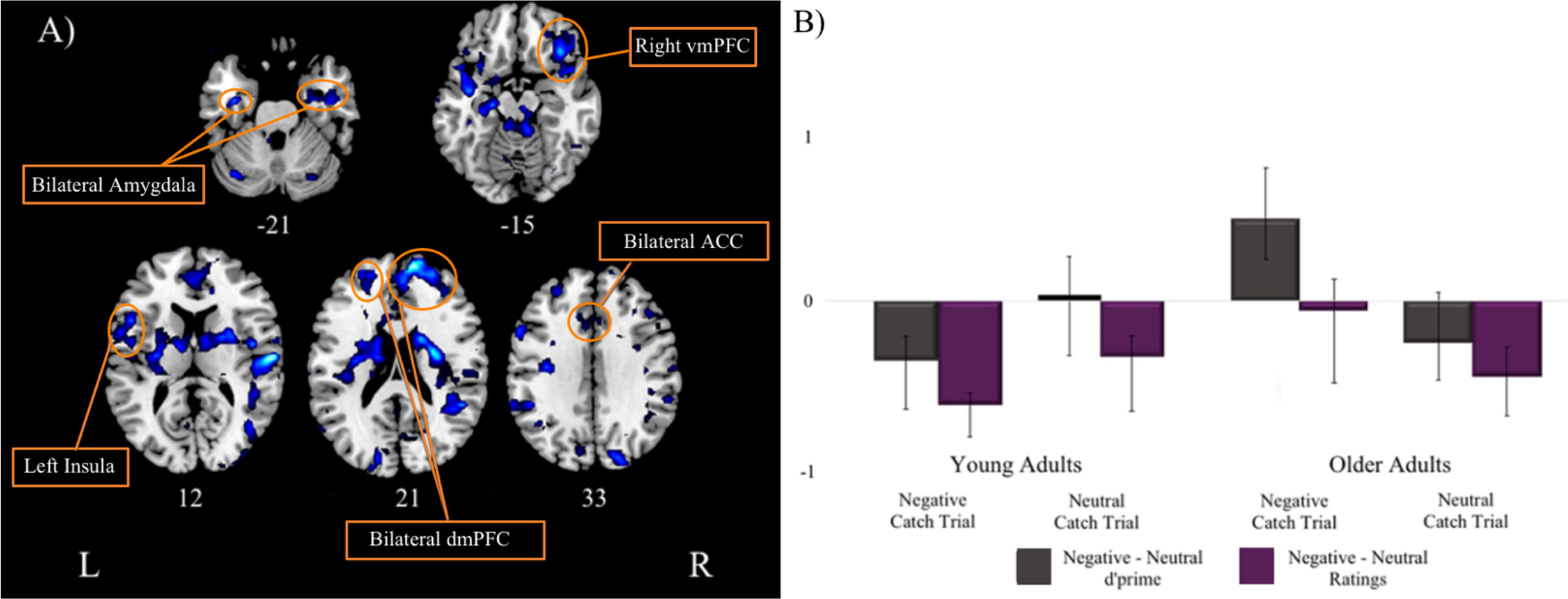
Singular image and correlation profile for the Behavioral PLS LV. A) Singular image for the behavioral PLS LV. Only negative salience regions (in blue) withstood our spatial extent cutoff (minimum of 15 clusters), threshold BSR of ±3.28, *p* < .001. B) Brain-behavior correlation profile for LV. The correlation profile indicates during negative catch trials, these negative salience regions are positively correlated with greater negative intensity ratings and with greater negative memory accuracy for young adults, but worse negative memory accuracy for older adults. During neutral catch trials, these negative salience regions are positively correlated with greater negative intensity ratings for both young and older adults. Error bars represent 95% confidence.

**Table 4.** Behavioral PLS LV brain regions in which task-related activity showed age effects and differentially related to memory accuracy and ratings.

| Temporal Lag | Bootstrap Ratio | Spatial Extent | MNI coordinates |  |  | HEM | Gyral Location | Brodmann Area |
| --- | --- | --- | --- | --- | --- | --- | --- | --- |
|  |  |  | x | y | z |  |  |  |
| Negative Salience Regions |  |  |  |  |  |  |  |  |
| 3 | -5.7118 | 28 | -36 | -9 | -27 | Left | Amygdala |  |
| 4,6 | -4.4107 | 51 | 24 | -3 | -27 | Right | Amygdala |  |
| 3 | -4.0841 | 15 | 18 | 66 | -3 | Right | Medial Frontal Gyrus (vmPFC) | 10 |
| 3 | -4.8688 | 15 | 3 | 39 | -21 | Right | Medial Frontal Gyrus (vmPFC) | 10/11 |
| 3,5,6 | -6.07 | 438 | -42 | 3 | -12 | Left | Insula* | 13 |
| 2,4 | -5.3362 | 145 | 33 | 30 | -6 | Right | Insula* | 13 |
| 2,3,4,5,6 | -5.356 | 253 | -15 | -87 | 24 | Left | Cuneus | 17/18 |
| 3,5 | -4.7151 | 38 | -12 | -51 | 0 | Left | Lingual Gyrus | 19 |
| 2,4,5 | -5.6808 | 238 | 42 | -69 | 15 | Right | Middle Occipital Gyrus | 19/39 |
| 3 | -4.4994 | 107 | 60 | -15 | 33 | Right | Postcentral Gyrus | 2/3 |
| 2 | -5.0489 | 15 | 15 | 36 | 12 | Right | Anterior Cingulate | 24/32 |
| 6 | -4.1266 | 35 | 0 | 3 | -6 | Left | Anterior Cingulate | 25 |
| 2,6 | -4.5913 | 40 | -6 | -15 | 45 | Left | Cingulate Gyrus | 31 |
| 2,3 | -4.7605 | 45 | 21 | -66 | 21 | Right | Posterior Cingulate | 31 |
| **3,5** | **-5.2796** | **111** | **-12** | **24** | **30** | **Left** | **Cingulate Gyrus (dACC)** | **24/32** |
| **2** | **-4.3596** | **51** | **3** | **21** | **33** | **Right** | **Cingulate Gyrus (dACC)** | **32** |
| 3,5,6 | -5.4848 | 119 | 57 | -69 | 3 | Right | Middle Temporal Gyrus | 21/37 |
| 2,3,5,6 | -6.6655 | 225 | -3 | 42 | 60 | Left | Superior Frontal Gyrus (premotor cortex) | 6 |
| 2,3,5,6 | -7.7168 | 27 | 45 | 12 | 54 | Right | Middle Frontal Gyrus (premotor cortex) | 6 |
| 3,4 | -5.392 | 15 | 15 | -57 | 42 | Right | Precuneus | 7 |
| 3,4 | -7.2164 | 553 | -27 | 42 | 45 | Left | Middle Frontal Gyrus*** | 8/9 |
| **2,4** | **-5.6301** | **92** | **-21** | **54** | **24** | **Left** | **Superior Frontal Gyrus (mPFC)** | **9/10** |
| **2,3,4,5,6** | **-7.1361** | **521** | **12** | **60** | **21** | **Right** | **Superior Frontal Gyrus (mPFC)** | **9/10** |
| 4 | -6.7549 | 1904 | -30 | -24 | -12 | Left | Hippocampus** |  |
| 2,6 | -5.0098 | 67 | -30 | -9 | 9 | Left | Putamen |  |
| 2,3,5 | -7.0149 | 2781 | 24 | 6 | 6 | Right | Putamen |  |

*Note:* HEM = hemisphere; \*extends into inferior frontal gyrus; \*\*extends into parahippocampus; \*\*\*extends into superior frontal gyrus; bolded regions are shown in Figure 3

#### 3.3.2 Cue Period Seed PLS

The Seed PLS analysis identified one significant LV. This LV, which accounted for 41.10% of the total cross-block covariance (*p* = 0.003) is presented in **Figure 5a**, and local maxima of selected regions for this LV are presented in **Table 5.** Only negative salience brain regions from this LV withstood our spatial extent cutoff of a minimum of 15 clusters and BSR threshold of ±3.28. The PLS correlation profile in **Figure 5b** demonstrates that activity in the right vmPFC seed is correlated with activity in this set of regions during negative and neutral catch trials for young adults. For older adults, activity in the right vmPFC seed is correlated with activity in this set of regions during neutral catch trials, but anticorrelated with activity in this set of regions during negative catch trials. These regions included: bilateral anterior cingulate, right amygdala, left hippocampus and the right ventrolateral prefrontal cortex.

**Fig. 5.**
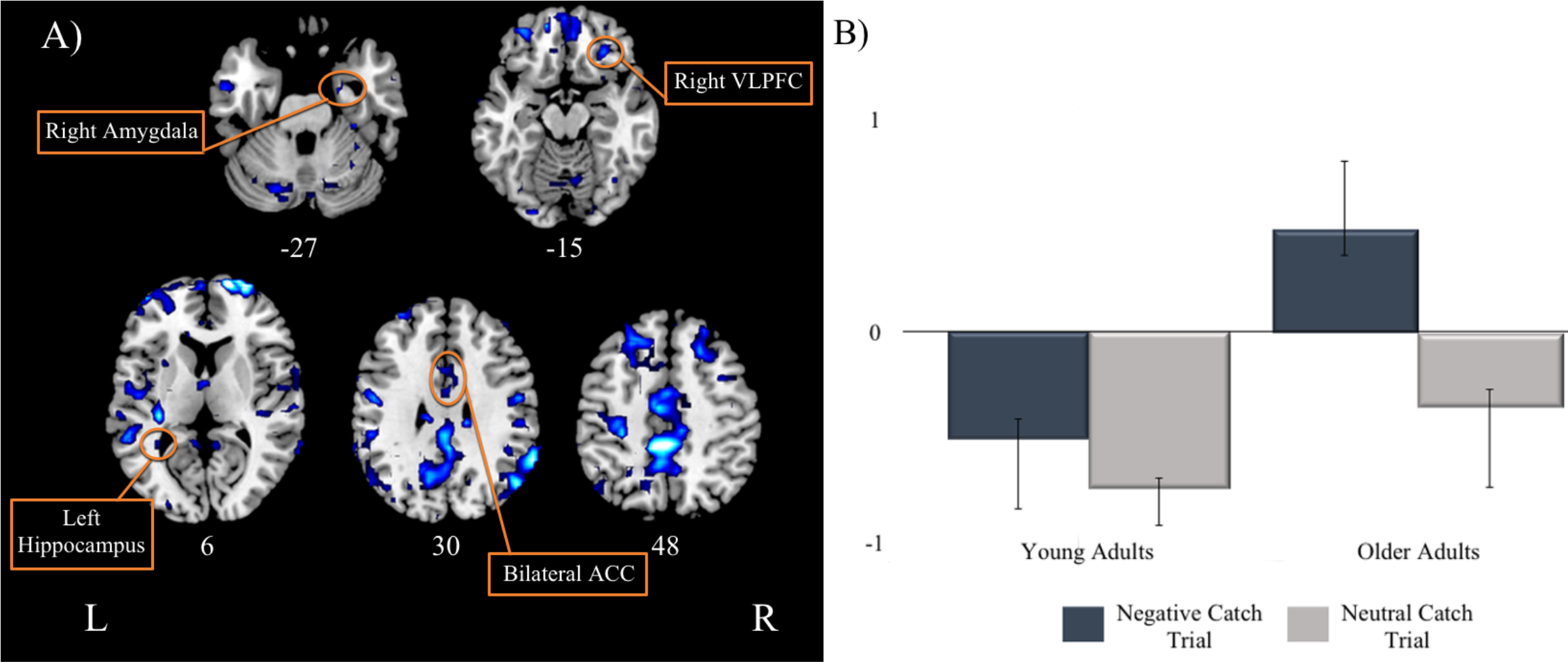
Singular image and correlation profile for the Seed PLS LV. A) Singular image for the seed PLS LV. Only negative salience regions (in blue) withstood our spatial extent cutoff (minimum of 15 clusters), threshold BSR of ±3.28, *p* < .001. B) Brain-behavior correlation profile for LV. The correlation profile indicates during negative and neutral catch trials for young adults and neutral catch trials for older adults, these negative salience regions are positively correlated with activity in the right vmPFC. During negative catch trials for older adults, these negative salience regions are anticorrelated with activity in the right vmPFC. Error bars represent 95% confidence.

**Table 5.**
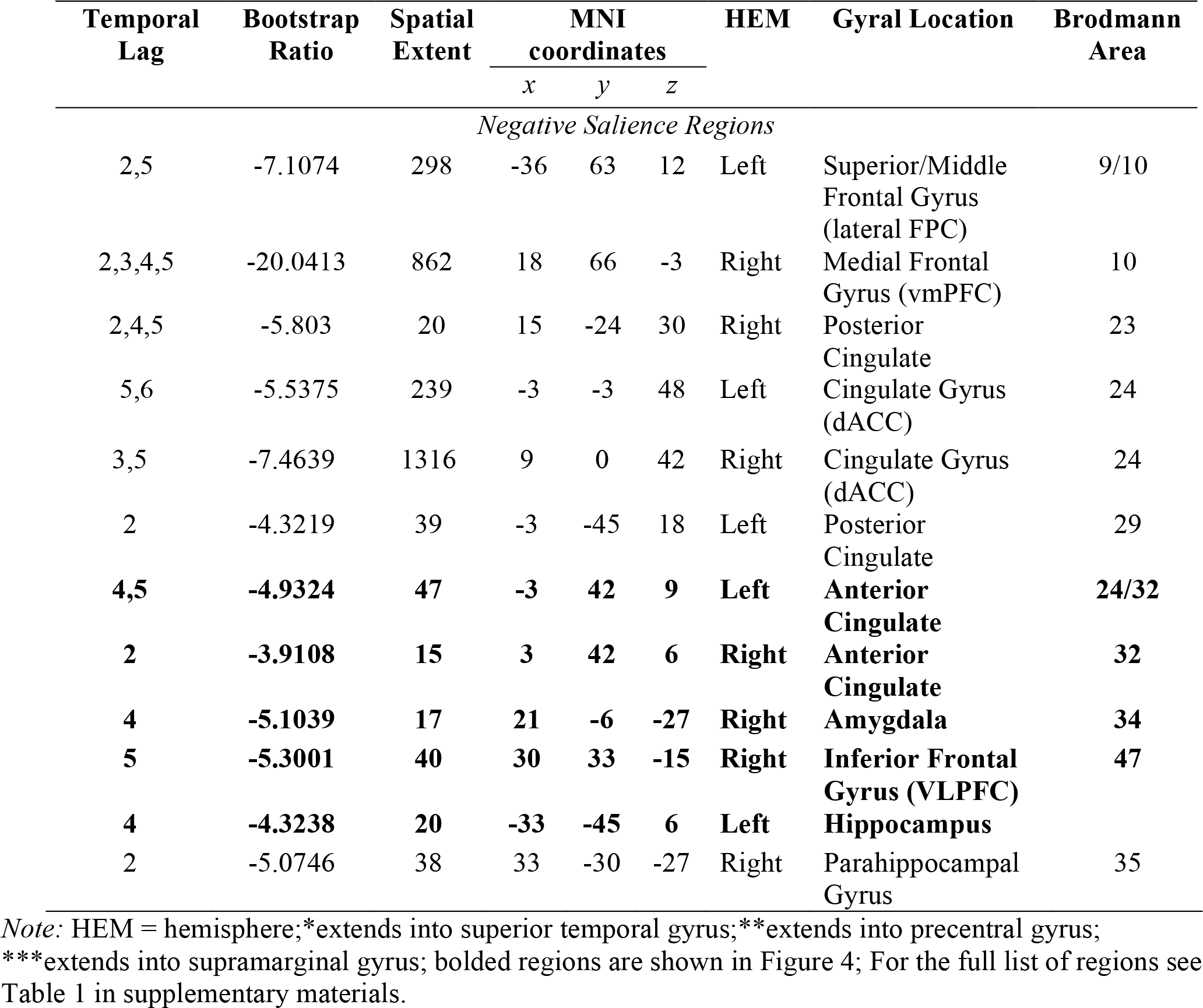
Regions functionally connected to the right vmPFC during negative and neutral catch trials for both young and older adults.

## 3. Discussion

The current study used event-related fMRI to investigate how anticipation affects episodic memory and the affective response associated with naturalistic emotional events in young and older adults. Emotional anticipation was induced by presenting participants with a cue that indicated if a negative or a neutral image was to follow, thereby allowing participants to engage in anticipatory processes. We additionally investigated the impact of emotional uncertainty on these anticipatory processes by violating participants’ expectations about the impending emotional stimuli. That is, a proportion of our cues invalidly indicated the valence of the impending image. We were able to isolate these anticipatory processes from the stimulus processes through our use of catch trials as described in the methods section. Utilizing PLS, we found that aging alters the manner in which brain regions supporting emotional processing are recruited in anticipation of emotional experiences. Consistent with previous findings, old but not young adults showed worse memory for negative compared to neutral events. Older adults recruit this set of regions to engage in spontaneous regulation following negative, but not neutral cues, whereas young adults recruit the same regions to engage in threat-related vigilance following both cues. This finding was further supported by the results of our seed PLS analysis in which older adults showed an anticorrelation between amygdala and ventromedial PFC activation following negative cues, indicative of downregulation of negative affect. The interpretation of these findings and their possible implications are discussed below.

### 3.1 Behavioral Results

Behaviorally, young adults remembered a similar number of negative and neutral images while older adults showed reduced memory for negative than neutral images. This was somewhat unexpected given that young adults typically remember more negative than neutral items (for review, Talmi et al., 2013). There are a couple of possible explanations for this discrepancy. First, we tested participants’ memory relatively shortly after encoding (~10-60 minutes). The ‘modulation hypothesis of emotional memory’ states that the emotional arousal of negative images induces the secretion of stress hormones that enhances activity in both the amygdala and medial temporal lobe (MTL), thereby modulating memory consolidation (Dolcos, LaBar, & Cabeza, 2004; McGaugh, 2004). Greater memory for negative images should only emerge after a more substantial delay (Cahill & McGaugh, 1998). It is likely that if we had delayed our recognition test to 48 hours or more, we would have seen this negative memory benefit. Another non-mutually exclusive explanation for this finding is that our validity manipulation resulted in encoding to take place under divided attention. Typically, the memory benefit for negative images is under conditions of full attention to the stimuli (for review, Talmi et al., 2013). Here, young adults were likely focusing, at least partially, on the cues to potentially determine which images were invalid versus valid, resulting in only a subset of their attention being allocated to stimulus valence (Schmidt & Saari, 2007; Talmi & McGarry, 2012). Consistent with this explanation is our finding that young adults remember more invalid than valid images, whereas older adults showed no difference. This likely reflects young adults, but not older adults sensitivity to the emotional uncertainty induced in our task.

In support of our hypothesis that older adults would spontaneously downregulate negative affect following cues, older adults remembered more neutral than negative images. This is consistent with the socioemotional selectivity theory (SST). Compared with young adults, older adults have been found to focus more on their emotional regulation goals, a consequence of which is attenuated processing of negative information (Carstensen et al., 1999; Mather & Carstensen, 2005). Also consistent with the SST was the lack of a difference in memory performance between valid and invalid images for older adults. It is possible that the 80/20 validity manipulation was insufficiently distracting to draw older adults’ attention away from emotion regulation, which they prioritize if permitted by task demands. Some other studies have found that when older adults have a challenging secondary task to perform, such as a concurrent auditory listening task while encoding emotional images, they perform similarly to the young, remembering more negative than neutral or positive images (Knight et al., 2007; Mather & Knight, 2005). This is consistent with the cognitive control model extension of SST that hypothesizes that emotion regulation is dependent on cognitive control (Mather & Knight, 2005).

In regard to our intensity ratings, we found that for both young and older adults, negative images, regardless of validity, induced a greater affective response than did neutral images. Interestingly, we found that both young and older adults rated negative valid images as more intense than negative invalid images. This is consistent with young adults engaging in threat-related vigilance due to their increased sensitivity to our cue-validity manipulation. This is inconsistent, however, with the idea that spontaneous downregulation during anticipation of negative events would manifest as reduced intensity ratings for valid than invalid stimuli. This discrepancy between affective ratings and memory has some precedent in the existing literature, however. Older adults are more likely to engage in suppression than any other type of emotional regulation strategy when not given instructions on how to regulate (Eldesouky & English, 2018; Nolen-Hoeksema & Aldao, 2011). Previous evidence suggests that suppression instructions generally result in impaired memory but has no effect on the subjective affective response (Gross, 2002). That is, affective response ratings for emotional stimuli are similar under passive viewing and directed suppression instructions, across age. Researchers have speculated that during suppression, young adults are focused on controlling their outward expression of negative affect (i.e. facial expression, vocal responses) at the expense of deep encoding of the emotional stimuli (for review, Dillon et al., 2007; Richards, 2004). In the current study, older adults may not have engaged in elaborative processing of the negative stimuli, at the expense of their memory. The imaging data discussed below support this. Further evidence supporting the notion that arousal ratings are not indicative of regulation in the current study is our finding that for both young and older adults, there was no correlation between intensity ratings and memory accuracy for negative or neutral images.

### 3.2 Older adults engage in spontaneous downregulation

Our behavioral PLS revealed a set of regions, including the medial PFC and amygdala, implicated in emotional processing and regulation that were differentially recruited with age. The vmPFC and dmPFC were revealed to be in the set of regions, both of which have been implicated in emotional and self-referential processing, with the vmPFC seeming to play a more crucial role in spontaneous downregulation (for further discussion, see Etkin, Egner, & Kalisch, 2011). The set of regions also included the amygdala, which is suggested to be involved in the detection and processing of emotional salience (Davis, 1992; Davis & Whalen, 2001), the insula, which is thought to play a more introspective role in emotional detection (Augustine, 1996; Critchley, Mathias, & Dolan, 2002; Phelps et al., 2001), the ACC, which is heavily involved in the generation of behavioral responses to emotional events (Craig, 2009; Critchley et al., 2002) and the hippocampus, which is implicated in episodic memory formation (for review, Yonelinas, 2013). For older adults, activity in these regions predicted worse negative memory accuracy relative to neutral memory accuracy following negative cues. We believe this pattern of results suggests that older adults engage in spontaneous downregulation of negative affect when they anticipate an impending negative image. The set of regions revealed is more consistent with those recruited during more automatic/spontaneous downregulation, namely suppression, than effortful/instructed downregulation, namely reappraisal. Specifically, suppression in older adults has been found to rely on the medial PFC, whereas reappraisal has been found to involve the lateral PFC (for further discussion see, Gyurak, Gross, & Etkin, 2011; and Phillips, Drevets, Rauch, & Lane, 2003). Collectively, our imaging and behavioral results add to the growing body of literature finding that older adults, but not young adults, engage in spontaneous suppression which results in impaired negative memory accuracy. Most notably though, this is the first study, to the best of our knowledge, to provide evidence that older adults engage in spontaneous suppression in anticipation, not just presentation, of negative events.

While behavioral PLS reveals the relationship between brain regions and behavioral measures, it does not reveal the relationship between the brain regions. We were particularly interested in the functional connectivity between the vmPFC and the amygdala for a few reasons. The vmPFC has been repeatedly indicated in emotional processing (Bush, Luu, & Posner, 2000; Ochsner et al., 2004; Taylor, Phan, Decker, & Liberzon, 2003), instructed down-regulation of negative affect (for review, Ochsner, 2008) and self-relevant processing (for review, Wagner, Haxby, & Heatherton, 2012). Neuroimaging evidence has shown that it is functionally connected with the amygdala (Sakaki et al., 2013; St Jacques et al., 2010; Urry et al., 2006). Furthermore, research in rodents and human lesion studies shows that the vmPFC has a direct anatomical connection to the amygdala. In rodents, direct high-frequency stimulation to the infralimbic cortex, the homolog to the vmPFC in humans, results in reduced responsiveness of neurons in the amygdala (Quirk, Likhtik, Pelletier, & Pare, 2003). Humans with bilateral vmPFC lesions show greater amygdala responses to negative images as well as elevated resting-state amygdala functional connectivity (Motzkin, Philippi, Wolf, Baskaya, & Koenigs, 2015). Collectively, these results suggest that connectivity between the vmPFC and amygdala may underlie older adults’ anticipatory downregulation of negative affect. Using seed PLS, we found that activity in the vmPFC was anticorrelated with activity in the amygdala, as well as the ACC, the hippocampus, the insula and visual association cortex areas following negative cues for older adults. By contrast, young adults showed positive connectivity in this same network. Previous perception and memory neuroimaging studies using naturalistic emotional events, like those used here, have demonstrated older adults, to a greater extent than young, show a negative coupling between vmPFC and amygdala activity (Gunning-Dixon et al., 2003; Leclerc & Kensinger, 2011; Roalf, Pruis, Stevens, & Janowsky, 2011; St Jacques et al., 2010). This negative connectivity pattern has been posited to represent older adults’ greater focus on emotional regulation goals that results in a top-down of the vmPFC on the amygdala. The current results extend previous findings by showing that older adults employ vmPFC-mediated emotional suppression strategies to reduce negative affect proactively, during anticipation of negative events. Notably, the negative connectivity between the vmPFC, the hippocampus, and visual association cortex areas further suggests that anticipatory emotion regulation engagement contributes to older adults’ impoverished negative memory accuracy through suppression of episodic encoding mechanisms.

It is important to mention that we cannot rule out the possibility that at least some of the results may be attributed to subjects, particularly older adults, averting their gaze to impending negative images. Previous eye tracking studies have found that older adults but not young adults, look away from negative stimuli, which they speculate may have resulted in their worse memory for negative stimuli (Isaacowitz et al., 2006; Knight et al., 2007). However, we do not believe that eye movements can fully account for the present pattern of results. First, if older adults were looking away when presented with negative cues, we would predict that their memory would be worse for neutral invalid (i.e. negatively cued) than neutral valid images and better for negative invalid (i.e. neutral cued) than negative valid images. Instead, we found that validity had no impact on memory performance for older adults. Further, it is unclear how the relationship we observed between activity in emotional processing regions and memory performance could be explained by older adults looking away from the screen when presented with a negative cue. Future fMRI studies incorporating eye tracking may prove useful in determining the presence and impact of eye movements on the neural processes of emotional regulation and its relationship with memory.

### 3.3 Young adults engage in threat-related vigilance

The set of emotional processing regions recruited during spontaneous downregulation by older adults was differently recruited by young adults. Specifically, activity in these regions predicted more intense ratings and better negative memory, following both negative and neutral cues in the young. This pattern of results is consistent with a threat-monitoring strategy in which young adults were constantly preparing for negative affect, regardless of the cue. As discussed above, young adults were particularly sensitive to the cue-validity manipulation, having better memory for invalid than valid images. This likely induced a sense of unpredictability, which has been shown previously to result in a threat-monitoring state in young adults (Bar-Anan et al., 2009; Grillon et al., 2004). This sense of unpredictability results in stronger affective responses (Bar-Anan et al., 2009; Grupe & Nitschke, 2011; Nader & Balleine, 2007; Yoshida et al., 2013) and similar engagement of emotional processing regions during negative and neutral events (Baas, Milstein, Donlevy, & Grillon, 2006; Nelson et al., 2015; Shackman, Maxwell, McMenamin, Greischar, & Davidson, 2011). Thus, our finding of activity in these regions predicting a stronger affective response following both negative and neutral cues is consistent with previous threat-monitoring research.

Further confirming that older and young adults differentially recruited the same emotional processing regions is the finding that activity in the vmPFC was positively correlated with activity in the amygdala, ACC, hippocampus, insula and visual association cortex areas following both negative and neutral cues. We observed the opposite pattern in older adults, in which the vmPFC was anticorrelated with activity in these regions. Rather than suppression, young adults recruited these regions together in a manner that is most consistent with hypervigilance for the detection of emotional salience (Baas et al., 2006; Nelson et al., 2015; Nitschke et al., 2006; Shackman et al., 2011).

It is important to describe the cues and their potential impact, distinct from their association with the stimuli. Negative and neutral cues depicted typical naturalistic sounds: screeching tires and wind chimes, respectively, which differed in arousal and valence, similarly to the stimuli they predicted. We selected these cues for several reasons. First, extensive piloting showed that young and older adults were equally able to remember the association between these cues and the stimulus valence they predicted. Older adults, in particular, had greater difficulty making cue-stimulus associations when cues were similarly neutral (i.e. X or O; different neutral tones), as have been used in some prior studies in young adults (Grupe et al., 2013; Mackiewicz et al., 2006; Nitschke et al., 2006; Sarinopoulos et al., 2010). Second, the use of auditory, rather than visual, cues eliminated the possibility that any anticipatory visual perceptual neural activity could be explained by perceptual processing of the cues. Lastly, the cues possess ecological validity in their association with neutral and negative emotional events. Nonetheless, one might argue that the cue-related activity that we attribute to emotional regulation, and age differences therein, reflects emotional processing of the cues rather than in anticipation of the stimuli, per se. We do not believe this is likely for a couple of reasons. First, both intensity ratings and memory performance differed as a function of cue validity, which can only be explained by participants using cues to anticipate the valence of the impending stimuli. Furthermore, cue-related neural activity was predictive of intensity and memory performance indices associated with the stimuli. If this activity was purely reflective of cue-related emotional reactivity, it is not clear why it would predict encoding relevant measures. Finally, it seems likely that any emotional response elicited by cues would habituate quickly as they repeat every trial. Indeed, we found no difference in emotional intensity ratings at the beginning and end of the experiment in a subsidiary analysis. Thus, we believe the current results are best explained as mobilization of emotional and memory related network activity in anticipation of impending emotional events.

## 4. Conclusion

The current results show that older adults, but not young adults, spontaneously engage in downregulation of negative affect, namely suppression during the anticipation of negative events. These novel results show that a contributor to the ‘positivity effect’ in aging is anticipatory suppression which leads to impaired memory for negative relative to neutral events. Future work should investigate how spontaneous suppression of negative emotion in older adults contributes to the positivity effect in real-world situations. One question that remains unanswered is when spontaneous suppression manifests in the adult lifespan. Future studies should include middle-aged adults to determine when this age-related shift to spontaneous suppression occurs.

## 5. Acknowledgements

We would like to thank our research participants and research assistants for their time and contribution to the study.

---

**Supplementary Table 1.** Regions functionally connected to the right vmPFC during negative and neutral catch trials for both young and older adults that were not presented in the main text.

| Temporal Lag | Bootstrap Ratio | Spatial Extent | MNI coordinates |  |  | HEM | Gyral Location | Brodmann Area |
| --- | --- | --- | --- | --- | --- | --- | --- | --- |
|  |  |  | x | y | z |  |  |  |
| Negative Salience Regions |  |  |  |  |  |  |  |  |
| 2,5 | -7.1074 | 298 | -36 | 63 | 12 | Left | Superior/Middle Frontal Gyrus (lateral FPC) | 9/10 |
| 2,3,4,5 | -20.0413 | 862 | 18 | 66 | -3 | Right | Medial Frontal Gyrus (vmPFC) | 10 |
| 3 | -8.6537 | 913 | -51 | -45 | 21 | Left | Superior Temporal Gyrus | 13 |
| 6 | -7.0083 | 97 | -27 | -96 | 12 | Left | Middle Occipital Gyrus | 18 |
| 6 | -4.125 | 40 | 21 | -84 | -6 | Right | Lingual Gyrus | 18 |
| 5,6 | -6.0907 | 44 | -24 | -90 | -12 | Left | Fusiform Gyrus | 19 |
| 5,6 | -5.6273 | 106 | 39 | -78 | 24 | Right | Middle Occipital Gyrus | 19 |
| 2,3,5 | -5.5567 | 24 | -54 | -30 | 0 | Left | Middle Temporal Gyrus* | 21/22 |
| 4 | -5.2219 | 43 | 57 | 3 | -18 | Right | Middle Temporal Gyrus | 21 |
| 3,6 | -5.1996 | 132 | 60 | -24 | 0 | Right | Superior Temporal Gyrus | 22/41 |
| 2,4,5 | -5.803 | 20 | 15 | -24 | 30 | Right | Posterior Cingulate | 23 |
| 5,6 | -5.5375 | 239 | -3 | -3 | 48 | Left | Cingulate Gyrus | 24 |
| 3,5 | -7.4639 | 1316 | 9 | 0 | 42 | Right | Cingulate Gyrus | 24 |
| 2 | -4.3219 | 39 | -3 | -45 | 18 | Left | Posterior Cingulate | 29 |
| 2,3,4,56 | -6.1681 | 33 | -42 | -18 | 66 | Left | Postcentral Gyrus** | 3/4 |
| 2,3,4,5,6 | -8.5096 | 3621 | -9 | -42 | 54 | Left | Precuneus | 7/31 |
| 4,5 | -4.9324 | 47 | -3 | 42 | 9 | Left | Anterior Cingulate | 24/32 |
| 2 | -3.9108 | 15 | 3 | 42 | 6 | Right | Anterior Cingulate | 32 |
| 4 | -5.1039 | 17 | 21 | -6 | -27 | Right | Amygdala | 34 |
| 2 | -5.0746 | 38 | 33 | -30 | -27 | Right | Parahippocampal Gyrus | 35 |
| 2,3,5 | -4.8374 | 46 | 63 | -60 | 12 | Right | Middle Temporal Gyrus*** | 39/40 |
| 5 | -5.3001 | 40 | 30 | 33 | -15 | Right | Inferior Frontal Gyrus | 47 |
| 2,6 | -6.0427 | 35 | 0 | -39 | 66 | Left | Paracentral Lobule | 5 |
| 2,4,5,6 | -5.9785 | 256 | -3 | 9 | 66 | Left | Superior/Medial Frontal Gyrus | 6 |
| 4,5,6 | -6.6602 | 82 | 39 | 9 | 54 | Right | Superior Frontal Gyrus | 6 |
| 2,4,5,6 | -7.258 | 85 | 30 | -51 | 63 | Right | Superior Parietal Lobule | 7 |
| 2,4 | -5.7311 | 23 | -12 | 57 | 42 | Left | Superior/Middle Frontal Gyrus (frontal eye field) | 8 |
| 2,3,4 | -4.9051 | 77 | 18 | 51 | 39 | Right | Superior/Middle Frontal Gyrus (frontal eye field) | 8 |
| 5 | -3.9815 | 15 | 51 | 42 | 21 | Right | Superior Frontal Gyrus | 9 |
| 2,4,5 | -5.7462 | 20 | -12 | -21 | 27 | Left | Caudate |  |
| **4** | **-4.3238** | **20** | **-33** | **-45** | **6** | **Left** | **Hippocampus** |  |

\*Regions in bold are regions presented in Table 5 in the main text.

